# Neutrophil Elastase Activates Macrophage MMPs, Promotes Cell Adhesion and Cytokine Production Via Integrin-Src Kinases Pathway

**DOI:** 10.1101/474395

**Authors:** Karina Krotova, Nazli Khodayari, Regina Oshins, George Aslanidi, Mark L. Brantly

## Abstract

There are a number of diseases characterized by the presence of neutrophil elastase (NE) activity in tissues including cystic fibrosis and alpha-1-antitrypsin deficiency induced lung destruction. It is generally accepted that NE actively contributes to this pathological process, but the precise mechanisms has yet to be determined. We hypothesized that NE activates the macrophages (M□) pro-inflammatory program. We demonstrate that following NE exposure, monocyte-derived M□ release proteolytic activity composed of several matrix metalloproteinases (MMPs) which could contribute to extracellular matrix (ECM) degradation. NE upregulates expression of M□ derived pro-inflammatory cytokines including TNFα, IL-1β, and IL-8. Thus, NE-activated M□ can contribute to tissue destruction through the proteolytic activity of metalloproteinases and by supporting chronic inflammation through expression of pro-inflammatory cytokines. We also demonstrate that NE increases M□ adhesion that is attenuated by antibodies specific to integrin subunits. We show that the effects of NE on M□ can be mediated through an activation of integrin pathways. In support of integrin involvement, we demonstrate that NE activates the Src kinase family, a hallmark of integrin signaling activation. Moreover, pretreatment of macrophages with a specific Src kinase inhibitor, PP2, completely prevents NE-induced inflammatory cytokine production. Taken together these findings indicate that NE has effect on lung destruction that extends beyond direct proteolytic degradation of matrix proteins.

## INTRODUCTION

Neutrophil elastase (NE) is a serine protease expressed by neutrophils and there are a number of pathological conditions associated with the presence of its activity in tissues (1–6). NE together with other proteins is stored in azurophil granules of neutrophils and is involved in anti-microbial defense. In response to infection or other inflammatory stimuli neutrophils quickly infiltrate organs and release the contents of granules into the extracellular space. Released NE participates in host defense against many pathogens (7–9), but needs to be neutralized in a timely to prevent destruction of host cells and extracellular matrix (ECM). The most known and well characterized pathological condition associated with unopposed NE is α1-antitrypsin (AAT) – deficiency induced emphysema. AAT, a glycoprotein released by liver in blood circulation, plays a major role in NE inactivation. Normal serum concentrations of AAT range from 20-53µM; in AAT-deficiency (AATD) the circulating AAT levels typically 5-fold less than normal as a consequence of several mutations in AAT gene (10–13).

Individuals with AATD have increased risk to develop pulmonary emphysema, a genetic form of chronic obstructive pulmonary disease (COPD), and the presence of unopposed NE is considered as a major factor responsible for lung degradation. In other forms of emphysema the role of NE is less obvious, but NE burden in lungs correlates with severity of disease (14,15). Additional experimental data suggest that NE not only directly destroys tissues, but also triggers a pro-inflammatory response in lung resident cells. Alveolar macrophages are critical cells that are activated (16,17), and lungs of individuals with COPD have increased number of M□ but not neutrophils (18). M□ not only support inflammation in lungs but also able to degrade extracellular matrix. Human studies and mice experiments highlight the importance of M□ released proteolytic activity in smoke-induced emphysema (19–23). In addition, recently published data suggest that M□ also play an important role in NE-induced emphysema, as depletion of macrophages in the lungs rescue mice from emphysema development (24).

In this work we analyzed the effect of NE exposure on activation of human monocyte-derived M□. We demonstrate that M□ exposure to NE leads to activation of several MMPs and increases expression of the pro-inflammatory cytokines IL-8, IL-1β, and TNFα. In addition, we show that NE promotes integrin-mediated adhesion of M□ and Src kinase activation. Moreover, cytokine production by M□ in response to NE treatment is mediated through Src kinases since specific Src inhibitor, PP2, abolishes the effects of NE on M□.

Based on these data we propose a mechanism of NE mediated activation of M□ through integrins and Src family kinases. Thus, it is possible that inhibition of this pathway could attenuate NE –induced inflammation and destruction in diseases characterized by the presence of free NE activity such as AATD.

## MATERIALS AND METHODS

### Reagents

All cell culture reagents unless specified elsewhere, were purchased from Life Technologies, (Carlsbad, CA, USA). All chemicals not specified are from Sigma-Aldrich (St. Louis, MO, USA). Fluorogenic peptide was purchased from R&D Systems (Minneapolis, MN, USA). Collagen Type I from rat tail (Corning), PP2 Src inhibitor – 2µM (Calbiochem).

### Cell culture

Primary peripheral blood mononuclear cells (PBMCs) were obtained from outpatient healthy volunteers (UF IRB protocol 08-2007). Monocyte-derived M□ were obtained from PBMCs plated in serum-free RPMI. Unattached lymphocytes were washed out after 1 h, and adherent monocytes were differentiated in M□ by culturing for 10 days in RPMI with 10% FBS with both GM-CSF (1 ng/Ml) and M-CSF (10 ng/Ml).

### MMP activity

MMP activity was assessed by a fluorogenic assay measuring 7-methoxycoumarin group (Mca) release from synthetic peptide Mca-PLGL-Dpa-AR-NH2 (RnD Systems, Minneapolis, MN, USA). After treatment with NE, media from M□ were collected, and aliquots of conditioned media were mixed with MMP assay buffer containing 10 µM fluorogenic peptide. Changes in fluorescence was monitored hourly at 320/405 nm.

### Zymography

Media collected from unstimulated or NE-stimulated M□ were analyzed by gelatin zymography. Samples were mixed with 2x Laemmli loading buffer without heating, and subjected to electrophoresis on 8% polyacrylamide gel containing 1 mg/ml gelatin. After gel electrophoresis was performed at 4°C, gels were incubated in 2.5% (v/v) Triton-X100 for 30 minutes to recover enzymatic activity, then overnight in 50 mM Tris (pH 8.0), 5 mM CaCl2, and 1µM ZnCl2 at 37°C. At the end of incubation gels were stained with 0.125% Coomassie Blue. The presence of MMPs appears as transparent bands on blue background.

### Gene expression measured by real-time PCR)RT-PCR)

RT-PCR was conducted using Applied Biosystems TaqMan commercial primers on an ABI Prism 7500 fast detection system using standard protocol; 18S mRNA was used as an internal reference. Quantification of relative gene expression was performed using the comparative threshold cycle (C_T_) method (25).

### Cytokine levels

IL-8 protein levels were measured in M□ conditioned media using Biolegend ELISA kit (Biolegend, San-Diego, CA) in range 15.6-1000 pg/ml. Protein levels of IL-1β and TNFα were assessed by using Luminex based kits (RnD systems, Minneapolis, MN) according to manufacture recommendations. The range of standard curve for IL-1β was 1.8-3,997 pg/ml, for TNFα - 1.3-2,144 pg/ml. Samples with cytokine levels in range of standard curve were analyzed.

### Flow cytometry

After treatment, M□ were recovered by gentle scraping of cells from the plate using cell lifter. Cell surface markers were analyzed by standard flow cytometry using a Beckman Galios and post-analyzed using Caluza software (Beckman Coulter, Indianapolis, IN). FITC-labeled anti-human CD14, CD11b, PE-Cy7 labeled anti-human CD206, APC-labeled anti-human HLA-DR, and corresponding isotype controls were purchased from Beckman Coulter (Indianapolis, IN). APC-labeled anti-human CD163, PE-Cy7 labeled anti-human CD11c, FITC-labeled anti-human CD44 and corresponding isotype controls were from BioLegend (San Diego, CA). Before staining, Fc receptors were blocked with human BD Fc Block, and 7-AAD (BD Biosciences, San Jose, CA) was added 5 min before analysis to discriminate dead cells.

### Adhesion assay

Wells in 96-well plate were coated with either 10 µg/ ml fibronectin or 50 µg/ ml Collagen I in PBS 50 µl/well for 1h at room temperature. Wells were then washed 2 times with PBS and free surfaces were blocked with 50 µg/ ml Collagen I or 0.5% BSA in PBS solution for 1h at room temperature. Before experiments, M□ were cultured overnight in serum-free media and/or treated with 10 ng/ml LPS. The next day cells were labeled with 5 µg/ml Calcein/AM in HBSS/Ca^2+^/Mg^2+^ buffer for 20 min, then washed several times and detached by treating with accutase and gentle scraping. Cells were spun-down, suspended in HBSS/Ca^2+^/Mg^2+^, and treated with 50 nM NE right before loading on coated plate, some samples were pretreated with blocking antibodies to CD11b (Biolegend, clone ICRF44), CD18 (Biolegend, clone TS1/18), CD29 (Biolegend, clone P5D2), CD11c (Biolegend, clone Bu15) in concentration 50 µg/ml 15 min on ice before adding NE. M□ were plated on pre-coated 96-well plate 100 µl per well in concentrations of 5×10^5^ cells/ml, and cells were left to adhere for 30 min at 37°C in a CO_2_ incubator. At the end of incubation fluorescence was read at 480/520 nm, with the intensity corresponding to total number of cells in wells. Then wells were washed with PBS four times to remove unbound cells, and fluorescence was measured again. The intensity of the second fluorescence measurement corresponded to the number of attached cells. To calculate the percentage of attached cells the fluorescence intensity from attached cells was divided by the fluorescence from all cells in corresponding well.

### Western blot analysis

M□ lysates were separated by 7.5% or 10% SDS-PAGE and transferred to a nitrocellulose membrane. Membranes were blotted with specific antibodies to phospho-Src (Cell Signaling), MMP-14 (EMD Millipore), GAPDH (Santa Cruz Biotechnology, Dallas, TX, USA).

### Statistics

All results are expressed as the mean ± S.E.M. Statistical analysis was performed using the two-tailed Student’s *t* test (GraphPad Prism 7.01 software; GraphPad Software, San Diego, CA, USA), and *p* < 0.05 was considered statistically significant. Data were plotted using GraphPad Prism 7.01.

## RESULTS

### Profile of MMPs activated by NE treatment of M□

To analyze the effect of NE on M□-released MMP activity, conditioned media was collected after stimulation with 50 nM NE for 18 h, and protease activity was measured by using MMP fluorescent substrate. Exposure to NE significantly increased proteolytic activity in M□ conditioned media which could be inhibited by 20 µM marimastat, a broad-range MMP-inhibitor, but not by 2 mM PMSF, a serine protease inhibitor (Fig. 1A). We did not find any changes in mRNA expression of any MMPs we studied in response to 50 nM NE (data not shown), though it has been shown that NE at higher concentration 166-500 nM can upregulate MMP-2 at transcriptional level (26). To estimate which MMPs contribute to elevated activity, we used gelatin zymography to detect proteases with gelatinase activity both in latent and in active form. Conditioned media from control cells had a major band around 92 kDa and minor band around 70 kDa corresponding to latent forms of MMP-9 and MMP-2 (Fig1B, Lane1). On the other hand, conditioned media from M□ stimulated with NE revealed several new bands on zymogram (Fig. 1B, lane 2 vs lane1); right under pro-MMP-9 band appeared new band of active MMP-9, under pro-MMP-2 - new band of active MMP-2, and also a ladder of several new bands around 60-40 kDa could be seen. It is known that NE can directly activate latent form of MMP-9 by proteolysis (27). It can also activate MMP-2, but indirectly with only the presence of cells which express MMP-14 on cell surface (28). In agreement, we found that treating conditioned media from M□ with NE leads to the appearance of the active form of MMP-9, however, the band corresponding to MMP-2 and bands around 60-40 kDa required the presence of cells during NE treatment as they did not appear if conditioned media was treated with NE (Fig. 1B, Lane 3). We hypothesized that one of the bands around 60-40 kDa could be MMP-14. Human M□ express MMP-14 and we found that NE targets MMP-14 causing it to shed from M□ cell surface in a concentration dependent manner (Fig. 1C). Incubation of gels in the presence of EDTA completely blocked the appearance of any bands which indicate they represent metal dependent enzymes such as MMP family (data not shown).

**FIGURE 1.**
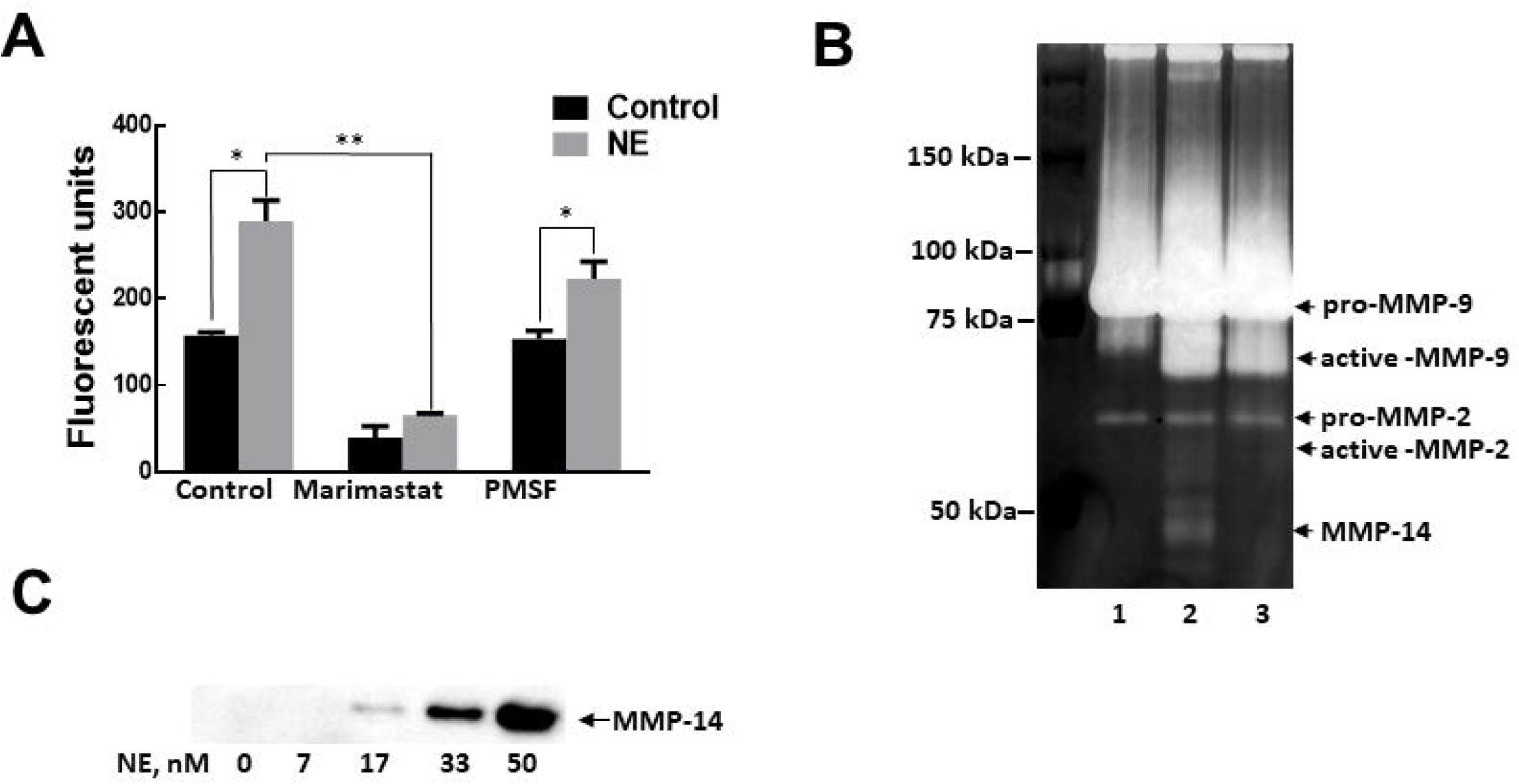
An increased M□ MMP activity after treatment with NE. **A.** M□ were treated with 50 nM NE for 18 h. The activity of MMPs was measured in M□ conditioned media by measuring fluorescence released from the digested peptide. To confirm that the digestion of substrate was MMP-specific, the reaction was also performed in the presence of 20 µM Marimastat or 2 mM PMSF. Data are shown from a representative of 3 independent experiments. \**p* < 0.05, \*\**p* < 0.01 **B.** Gelatin zymography of M□ conditioned media. In lane 1 - media from control cells. Lane 2 – media from M□ treated with 50 nM NE for 18h. Lane 3 – 50 nM NE was added to conditioned media from control cells. Representative image of 4 independent experiments **C.** Immunoblot of MMP-14 present in conditioned media of macrophages after treatment with different concentrations of NE for 18 h. Representative image of 3 independent experiments with similar results.

### NE upregulates inflammatory cytokines production in M□

Next we analyzed changes in cytokine levels expressed by M□ in the presence of NE. We found that 50 nM NE significantly increased IL-8 levels released by M□ (Fig. 2A and upregulation of IL-8 mRNA (Fig. 2B). In addition, NE induced IL-1β, and TNFα on mRNA levels, however, protein levels were below the limit of detection. In order to determine if NE affects the levels of these cytokines in M□, cells were co-stimulated with LPS and NE. We found that NE significantly enhanced LPS-induced IL-1β and TNFα cytokine production, though the response greatly varied between M□ derived from different individuals (Fig. 2C). However, NE did not add-up to IL-8 levels already increased by LPS. Increased protein levels for IL-1β were reflected by IL-1β mRNA upregulation, however for TNFα changes in mRNA levels did not reach statistical significance (Fig. 2D). The discrepancy in TNFα protein and mRNA levels likely resulted from the design of experiment when conditioned media for cytokine levels and cells for RNA isolation were collected at the same time, after 18h of treatment. While prolonged time treatment is optimal to see the accumulation of protein in media, this time point is not good for analysis of TNFα mRNA as it has been shown that in response to LPS treatment, TNFα mRNA expression increased within the first two hours and then dramatically declined (29).

**FIGURE 2.**
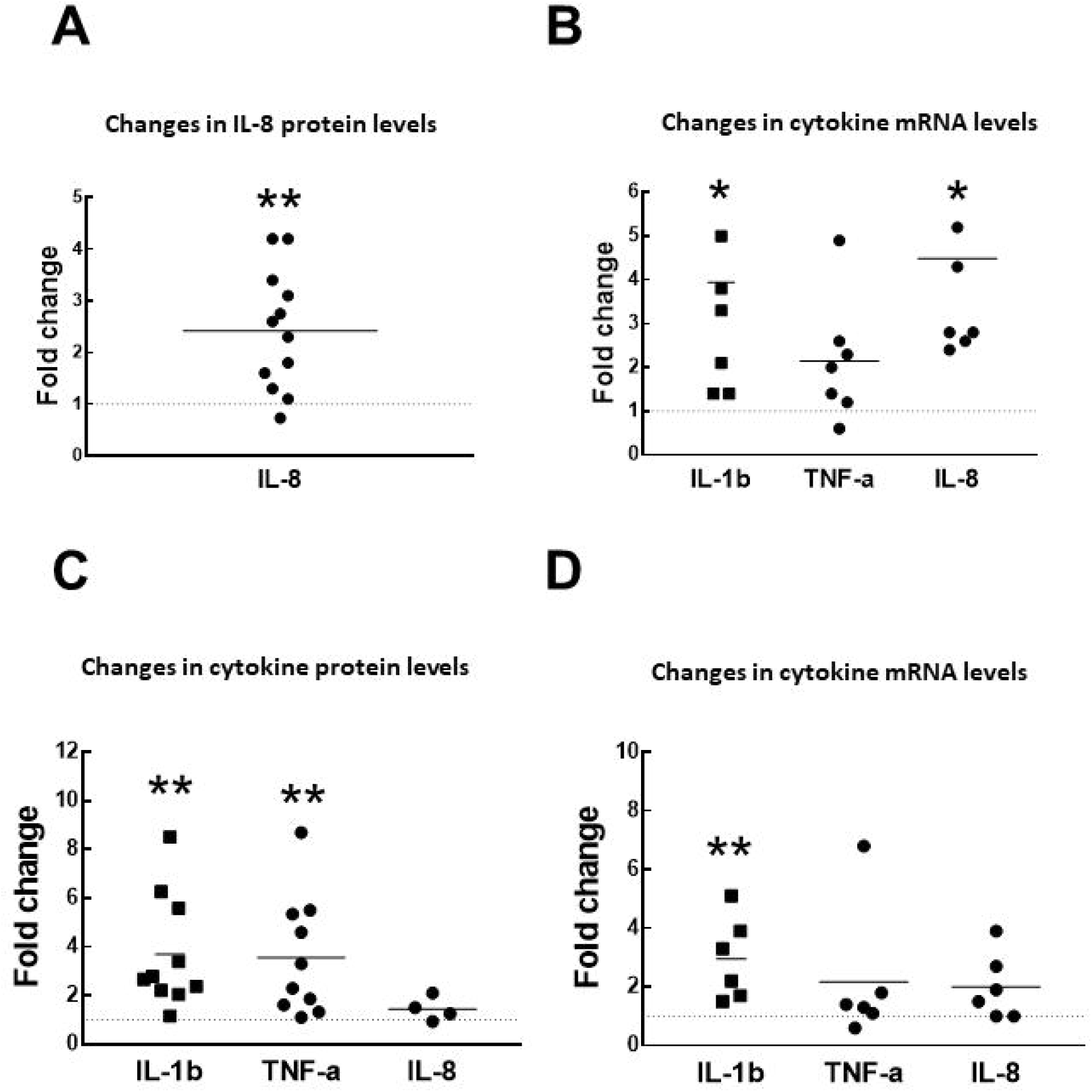
NE upregulates M□ cytokine levels. **A. and B.** M□ were treated with 50 nM NE for 18h. Protein levels for IL-8 were measured in conditioned media (A). mRNA levels for IL-8, TNFα, and IL-1β were measured by real-time PCR. (B) **C. and D.** M□ were treated with 50 nM NE and 10 ng/Ml LPS for 18 hrs. Control cells were treated with LPS only. Cytokine levels were measured in conditioned media (**C**). mRNA for cytokines was measured by real-time PCR (**D**). It should be noted that the set of data for IL-8 levels in C is limited, since a number of values was too high and out of range, and was not included in analysis. Each data point represents a single individual. Value 1 corresponds to no changes is marked on graphs by the horizontal dotted line. Data are presented as fold of changes in NE treated vs control cells. t-test for Control vs NE treated: \**p* < 0.05, \*\**p* < 0.01

### NE increases M□ adhesion

We observed that M□ treated with 50 nM NE overnight significantly changed their morphology. Treated M□ are more spread out on culture dish surface compare to non-treated cells (Fig. 3A). Calculations showed the trend in increase of area occupied by cells treated with NE but it did not reach statistical significance (Fig. 3B). The observation appeared to indicate enhanced M□ adhesion in the presence of NE. To test this, we analyzed how NE affect the attachment of freshly seeded M□ by using adhesion assay. At first, we planned to measure the adhesion of M□ to plates coated with fibronectin or collagen I. However, we found that collagen I was very poor matrix for M□ adhesion, much worse than BSA which is usually used in adhesion assays to block free space on plate surface. Only 2.9% ± 0.4% of M□ were bound to Collagen I-coated surface, while fibronectin-coated plates bound 29.3%± 7.3%, and BSA –coated plates bound 21.9% ± 1.6% of M□ loaded to plate (Fig. 4A). These preliminary data indicated that BSA is a good binding matrix for M□, and hence, is not suitable as a blocking agent. Since we found that collagen I had a low binding capacity for M□ in our experiments, we use it as a blocking agent after coating plates with fibronectin. Indeed, as seen in Fig. 4B collagen I significantly reduced background levels, and the difference between fibronectin-coated and non-coated wells was more dramatic with collagen I as a blocking reagent compared to BSA (11.8% vs 1.1% with collagen, 52.4% vs 43.8% with BSA). Therefore, in our experiments with NE we used collagen I as a blocking solution. Plating M□ on fibronectin coated plates in the presence of 50 nM NE dramatically increased the number of attached cells (from 11.6 ± 1.1% in control to 24.1 ± 0.6 % in NE treated). Integrins are primary receptors which regulate cell adhesion to extracellular matrix (30), hence, the increased M□ spreading might result from integrin activation. In agreement, we found that the pre-incubation of M□ with integrin antibodies specific to CD11b, CD18, or CD29 prevented enhanced binding to fibronectin under NE treatment (Fig. 4C). Visualization of Calcein-stained M□ used in adhesion assays under microscope confirmed that more cells were attached to the plate and also they were more spreading on plate surface compare to control cells (Fig. 4D).

**FIGURE 3.**
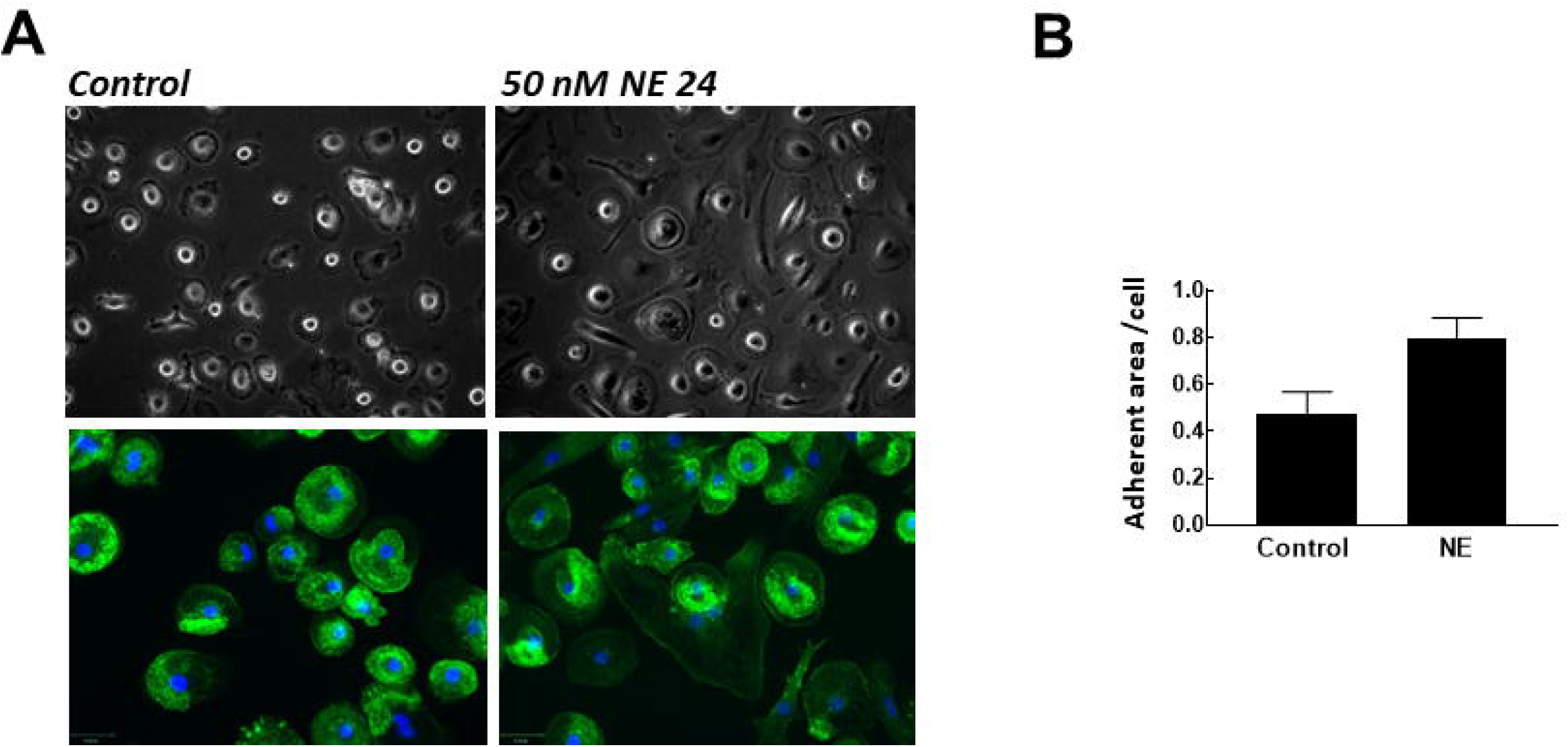
NE cause M□ spreading. **A.** Top panel. Phase contrast image of M□ after incubation with 50 nM NE for 24 h. Bottom panel. After treatment with 50 nM NE for 24 h, cells were fixed and F-actin was stained with FITC-phalloidin and nuclei were stained with DAPI. M□ look bigger compare to non-treated cells. **B.** Quantification of area under the cells which was expressed as a percentage from total view area. Data are representing of 3 independent experiments.

**FIGURE 4.**
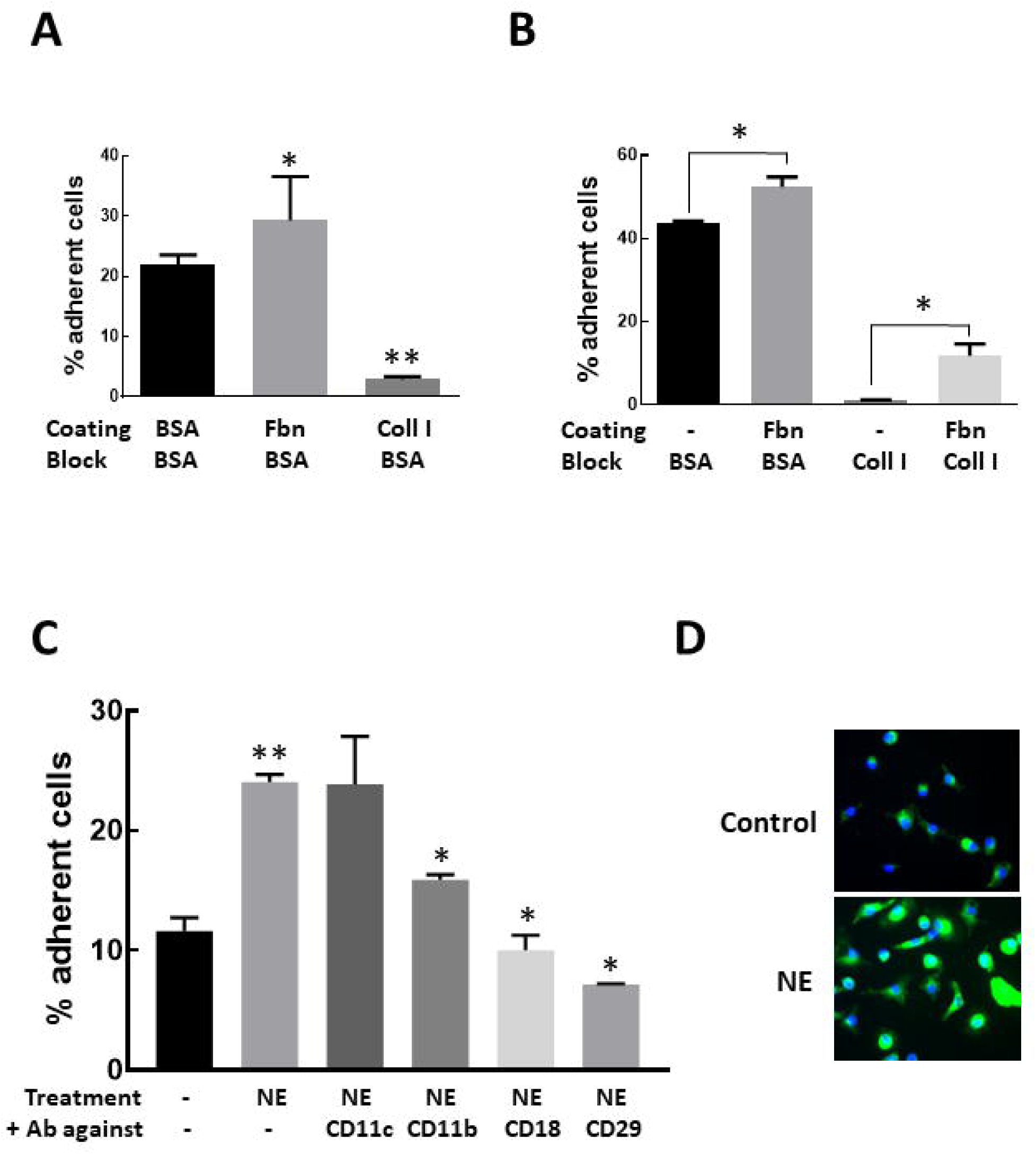
NE induces increased M□ adhesion **A.** and **B.** Adhesion experiments set-up. **A.** Adhesion of non-treated M□ to different matrix \**p* < 0.05 vs BSA coated, \*\**p* < 0.01 vs BSA coated **B.** Comparison of BSA and Collagen I as a blocking reagent \**p* < 0.05 **C.** and **D.** NE induced M□ adhesion to fibronectin (Fbn) is integrin-dependent **C.** The attachment of M□ to fibronectin coated – plates in the presence of NE, with or without the pretreatment of M□ with integrin-specific antibodies. \**p* < 0.01 vs NE treated, \*\**p* < 0.01 vs Control **D.** Representative images of Calcein-AM and Hoechst stained M□ confirmed that greater number of M□ adhere to fibronectin under NE treatment. All data are representatives at least three independent experiments.

### NE selectively cleaves some receptors but not integrins from the surface of M□

It has been shown that NE mediates many of its effects through cleavage of a number of cell surface molecules (1). This is especially true when NE was used in µM range concentrations (31,32), however, the effect of lower nM concentrations could be different. We were interested in how NE treatment changes the levels of integrin subunits and other M□ cell surface markers. In our next experiments we treated M□ with 200 nM NE, the concentrations similar to that found in lungs of AATD patients (33). In addition to integrin molecules CD11b and CD11c, we analyzed the effect of NE on cell surface markers we used to characterize PBMC-derived M□: CD163, CD44, HLA-DR, CD14, and CD206. We found that NE treatment did not change the levels of integrin subunits CD11b and CD11c. In contrast, NE caused significant loss of CD14 (54.1%±25.1%), CD44 (68.8%±7.1%) and CD206 (57.8%±20.2%). The changes in HLA-DR and CD163 levels in response to NE were highly variable between M□ derived from different individuals, but overall were not statistically significant (Fig. 5).

**FIGURE 5.**
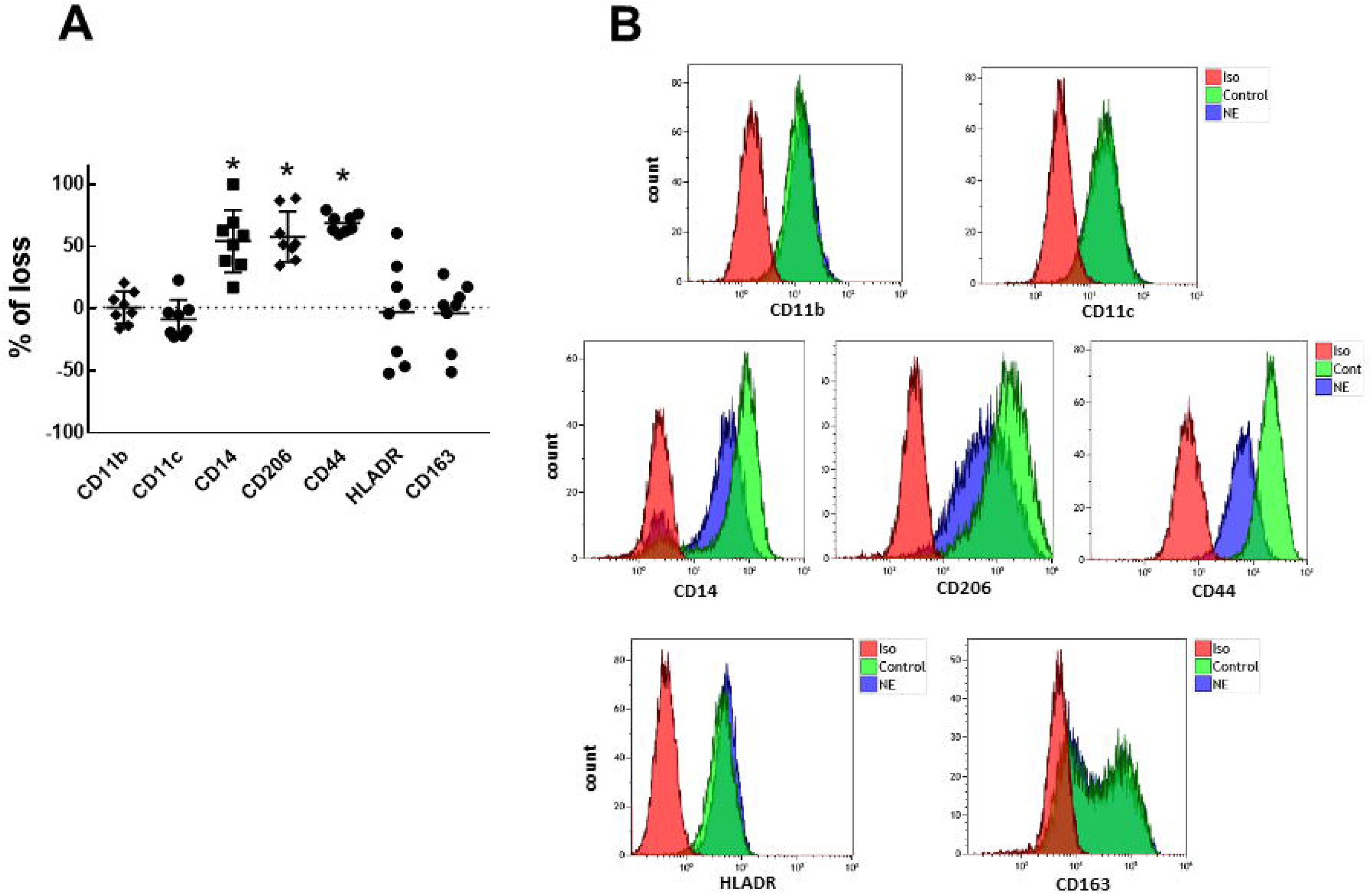
NE cleaves selected M□ surface receptors. **A.** M□ derived from 8 different individuals were treated with 200 nM NE for 1h and were analyzed by flow cytometry. The data presented as the percent of change in geometric mean of fluorescence intensity (MFI) for markers in NE treated cells relative to non-treated, control cells \**p* < 0.01 vs control **B.** Representative images of flow data summarized in A.

### NE-induces Src phosphorylation and Src inhibitor prevents upregulation of cytokines

Integrins participate in many cellular functions of immune cells through recruitment and activation of signaling proteins such as the members of Src non-receptor tyrosine kinase family (34–36). We determined if Src kinases were involved in signaling following M□ exposure to NE. The activation of Src was estimated using antibodies specific to phosphorylated tyrosine residue 419, which is autophosphorylated upon activation (37). We found that NE induces phosphorylation of Src family members as early as 10 min after stimulation with NE, with maximum phosphorylation at 30 min (Fig. 6A). The antibody we used were not isoform specific and recognized 3 bands in macrophage lysates indicating that at least 3 members of Src family kinases respond to NE treatment by phosphorylation. To confirm the involvement of Src family kinases in NE signaling, we pre-treated M□ with 2 µM of the specific Src inhibitor, PP2, before stimulation with NE. PP2 completely abolished NE-induced phosphorylation of Src (Fig. 6 B). In addition, PP2 caused a reduction in IL-8 and IL-1β expression both on mRNA and protein levels (Fig. 6 C and D), indicating that NE upregulates these cytokines through Src kinases.

**FIGURE 6.**
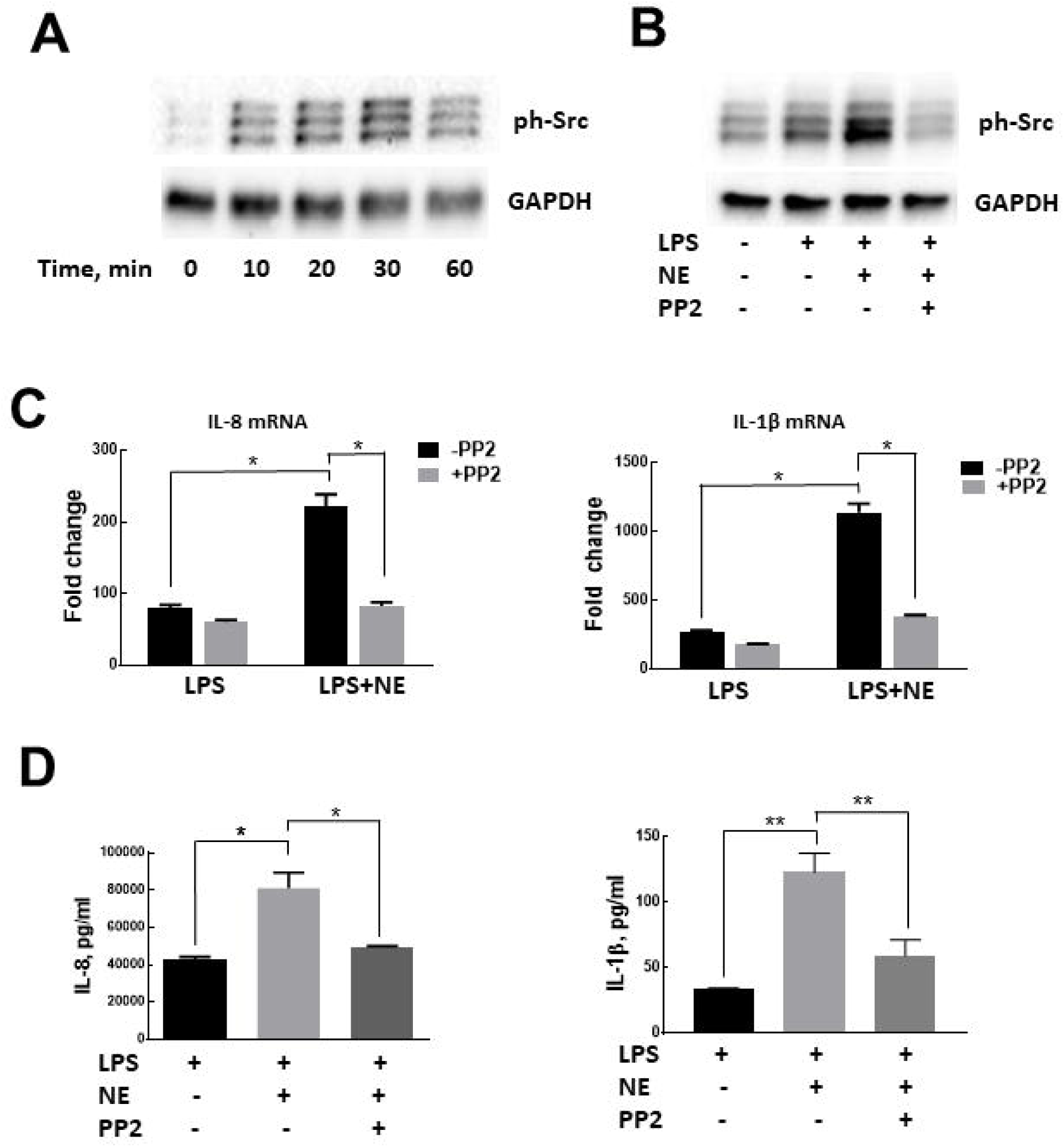
NE activates cytokine production through Src kinases. **A.** Time dependent phosphorylation of Src kinases in response to NE treatment. **B.** Pretreatment of M□ with PP2 abolished Src phosphorylation by NE treatment. Representative images of 3 independent experiments. **C.** The treatment with 2µM PP2 blocked NE-induced IL-8 and IL-1β cytokines production both on mRNA (**C**) and protein (**D**) levels. \**p* < 0.01, \*\**p* < 0.05.

## DISCUSSION

The purpose of this study was to show that NE is able to directly activate proteolytic and inflammatory activity of human M□ thus contributing to tissue destruction during lung diseases such as AATD-induced emphysema and COPD.

In our experiments we used concentrations of NE in nM range which correspond to NE levels in lungs of individuals with COPD (33,38,39). First, we showed that the incubation of human M□ with NE leads to significant increase in released MMP activity. This activity was composed of several MMPs including MMP-9, MMP-2, and MMP-14. In addition, we found that NE effects on MMP activities in M□ conditioned media were mediated through different mechanisms. MMP-9 and MMP-2 were released by M□ in extracellular space in latent forms, where MMP-9 was activated by NE directly through proteolysis. MMP-2 was activated by NE indirectly, only in the presence of M□, and MMP-14 which is a transmembrane protein was shed from the M□ cell surface (Fig. 1). While activation of MMP-9 and MMP-2 by NE was previously shown (27,28), this study demonstrates the shedding of active MMP-14 from cell surfaces following exposure to NE. Soluble forms of MMP-14 with retained activity have been detected in human bronchial asthma and several other pathologic conditions but mechanism of shedding was unknown (40,41). We identify here that the exposure of cells to free NE is a possible mechanism contributing to the presence of soluble forms of MMP-14. Hence, these data indicate that NE not only directly degrades ECM but also activates a number of MMPs which could amplify the matrix degradation.

Another way M□ are involved in tissue destruction is their support of chronic inflammatory conditions. Hence, we analyzed the effect of NE on the inflammatory cytokines expression in M□. We found that 50 nM NE induced a several fold increase in IL-8 mRNA and protein production by M□. IL-8, a potent neutrophil chemoattractant, implicated in pathogenesis of lung diseases such as COPD and cystic fibrosis (42,43). Moreover, a positive correlation between the levels of NE and IL-8 has been shown for COPD patients (38), however the source of IL-8 production was not determined. The two-fold increase in IL-8 production under exposure to NE was also reported for airway epithelial cells (44–46). Hence, in lungs both epithelial cells and M□ can contribute to elevated levels of IL-8 in the presence of NE. In addition to stimulation of IL-8 production, the presence of NE significantly augmented LPS-induced IL-1β and TNFα production (Fig. 2C and D). The upregulation of cytokines on transcriptional level we observed are consistent with a number of publications which demonstrate stimulatory effects of NE on cytokine levels both in mouse models and different types of human cells (44–49). It should be noted that downregulation of cytokines through degradation by NE was also reported, but with much higher NE doses than used in the current study (32,39,50–52). In summary, our data indicate that NE in concentrations found in COPD patients upregulates pro-inflammatory cytokines produced by human M□ that contribute to chronic inflammatory environment in lung, a hallmark of COPD (53).

A number of publications points out TLR-4 receptor and NF-kB transcriptional factor as mediators involved in stimulation of cytokines expression by NE (44,47). In this study we identified another pathway through which NE induced cytokine expression. We show that in M□ NE activate integrins and integrin-mediated intracellular signaling. First, we demonstrate that NE treatment dramatically increases M□ spreading and adhesion and this effect can be abolished by antibodies specific to integrin’s subunits CD11b, CD29, and CD18. The activation of Src kinases in response to NE stimulation further supported that NE targets integrins and integrin-mediated intracellular signaling. Our data complement previously identified signaling pathways triggered by NE since Src kinase activation is an early response element in integrin signaling which leads to complex cell signaling cross-talk including NF-kB activation (34). It is interesting to mention that the mechanism of NE-initiated inflammation via TLR-4 is associated with decreased TLR-4 surface expression (46). In contrast, we found that though integrins involved in NE signaling, their levels on M□ cell surface were not affected by NE treatment. At the same time exposure to NE led to loss of CD44, another adhesion molecule abundantly expressed on M□. CD44 is a hyaluronic acid receptor, but also exhibits affinity to numerous factors, e.g. collagen, fibronectin, chondroitin sulfate, osteopontin, and others (54). Both integrins and CD44 are used by macrophages for adhesion and migration, and implicated in different pathologic conditions. Our data indicate that NE treatment changes the balance of adhesion receptors in favor of integrins, however, the physiological significance of these findings is unclear and requires additional experiments. The activation of integrin signaling by NE seems not limited to M□, and involvement of NE, as well as another serine protease, cathepsin G in activation of integrin signaling and MIP-2 secretion has been reported for neutrophils(55).

Our study shows that NE increases integrin-mediated adhesion of M□ and modulate cytokine release from M□ through Src activation since Src kinase inhibitor PP2 inhibited the increase in cytokine expression. Recently, activated Hck (hematopoietic cell kinase), a myeloid-specific Src family kinase, was implied in lung –associated diseases (56,57). It has been shown that mice with a constitutively active Hck mutant develop in the lungs areas of mild emphysema and fibrosis (58). Our work proposes the mechanism of Src kinase activation through the presence of free NE, however the link between free NE and Src kinase activation needs to be further evaluated in in vivo experiments.

In conclusion our data highlight that NE can activate multiple pathways in M□ which can contribute to lung tissue destruction. In addition to the lung, free NE was detected in other tissues in association with pathological conditions (2–5). Our data elucidate mechanism of NE action and provide rationale for use of NE inhibitors as well as inhibitors of down-stream pathways such as Src kinases in diseases associated with NE-induced inflammation.

